# Neuroendocrine mechanisms governing sex-differences in chronic pain involve prolactin receptor sensory neuron signaling

**DOI:** 10.1101/2020.04.25.061663

**Authors:** Candler Paige, Priscilla A. Barba-Escobedo, Jennifer Mecklenburg, Mayur Patil, Vincent Goffin, David Grattan, Gregory Dussor, Armen N. Akopian, Theodore J. Price

## Abstract

Many clinical and preclinical studies report higher prevalence and severity of chronic pain in females. We used hyperalgesic priming with interleukin 6 (IL-6) priming and PGE_2_ as a second stimulus as a model for pain chronicity. Intraplantar IL-6 induced hypersensitivity was similar in magnitude and duration in both males and females, while both paw and intrathecal PGE_2_ hypersensitivity was more persistent in females. This difference in PGE_2_ response was dependent on both circulating estrogen and translation regulation signaling in the spinal cord. In males, the duration of hypersensitivity was regulated by testosterone. Since the prolactin receptor (Prlr) is regulated by reproductive hormones and is female-selectively activated in sensory neurons, we evaluated whether Prlr signaling contributes to hyperalgesic priming. Using ΔPRL, a competitive Prlr antagonist, and a mouse line with ablated Prlr in the Nav1.8 sensory neuronal population, we show that Prlr in sensory neurons is necessary for the development of hyperalgesic priming in female but not male mice. Overall, sex-specific mechanisms in the initiation and maintenance of chronic pain are regulated by the neuroendocrine system and, specifically, sensory neuronal Prlr signaling.

**Significance Statement:** Females are more likely to experience chronic pain than males, but the mechanisms that underlie this sex difference are not completely understood. Here, we demonstrate that the duration of mechanical hypersensitivity is dependent on circulating sex hormones in mice – where estrogen caused an extension of sensitivity and testosterone was responsible for a decrease in the duration of the hyperalgesic priming model of chronic pain. Additionally, we demonstrated that Prolactin receptor expression in Nav1.8^+^ neurons was necessary for hyperalgesic priming in female, but not male mice. Our work demonstrates a female-specific mechanism for the promotion of chronic pain involving the neuroendrocrine system and mediated by sensory neuronal prolactin receptor.

## Introduction

Many chronic pain conditions, such as migraine, fibromyalgia, temporomandibular joint disorders (TMD/TMJ), irritable bowel syndrome (IBS), and rheumatoid arthritis have a 2 to 6-fold greater prevalence or symptom severity in women when compared to men (Unruh, 1996; Berkley, 1997; Fillingim et al., 2009; Traub and Ji, 2013). Pain symptoms in women with these chronic pain conditions can change during the menstrual cycle as gonadal hormone concentrations fluctuate, and some, but not all, pain conditions decrease in frequency or intensity after menopause (Houghton et al., 2002; LeResche et al., 2003; Slade et al., 2011; Mathew et al., 2013). There is a general consensus that plasticity mechanisms in peripheral and central nociceptive pathways are critical for chronic pain development in males and females, but the precise mechanisms governing this plasticity are increasingly recognized as sex dimorphic and are still largely unknown. Nevertheless, recent progress was made in understanding underlying mechanisms for sex-dependent mechanisms of nociceptive plasticity (Mogil et al., 2011; Sorge et al., 2011; Sorge et al., 2015; Rosen et al., 2017; Martin et al., 2019). These findings on sex differences in nociceptive plasticity mechanisms, combined with abundant clinical and rodent data on the effects of gonadal hormones on pain, indicate a critical role for gonadal hormones in regulation of pain chronification (Fillingim et al., 2009; Traub and Ji, 2013).

Clearly there are gonadal hormone-regulated mechanisms that promote chronic pain in females, but these mechanisms have not been thoroughly characterized. Prolactin (PRL) and its receptor (Prlr) are prime candidates for this potential mechanism, since responsiveness to PRL in a variety of cells, including sensory neurons, is closely regulated by estrogen (Childs et al., 1999; Pi and Voogt, 2002; Diogenes et al., 2006; Belugin et al., 2013). Thus, Prlr signaling sensitizes pain-related ion channels and causes increased excitability in nociceptors specifically in females (Diogenes et al., 2006; Patil et al., 2013b; Liu et al., 2016; Patil et al., 2019b; Patil et al., 2019a). Moreover, Prlr function in female nociceptors is governed by estrogen signaling via a non-genomic pathway that involves sex-specific translation of Prlr mRNA (Patil et al., 2019b). Based on these previous studies, we hypothesized that Prlr signaling might differentially contribute to female-specific regulation of chronic pain that can be assessed by use of the hyperalgesic priming model (Aley et al., 2000). Our work reveals that initiation, maintenance and magnitude of hyperalgesic priming is governed by estrogen-dependent regulation of Prlr signaling in sensory neurons. Therefore, PRL signaling to Prlr is a gonadal hormone-dependent mechanism that promotes plasticity in the nociceptive pathway supporting development of chronic pain specifically in females.

## Materials and Methods

### Animals

All animal experiments were approved by the University Texas Health Science Center at San Antonio (UTHSCSA) and University of Texas at Dallas (UTD) Institutional Animal Care and Use Committee (IACUC). We followed guidelines issued by the National Institutes of Health (NIH) and the Society for Neuroscience (SfN) to minimize suffering and the number of animals used.

### Key reagents and mouse lines

8-12 week old female and male mice were purchased from Jackson Laboratory (Bar Harbor, ME). Ovariectomized (OVX) and gonadectomized (GdX) mice were purchased from Jackson Laboratory (Bar Harbor, ME). The estrous phases in adult females were determine by vaginal gavage as described by Caligionin (Caligioni, 2009). Estrogen and testosterone replacement procedures to generate OVX-E-2 and GdX-T mice were performed as previously described (Diogenes et al., 2006; Nettleship et al., 2007). 17β-estradiol (E-2; 300 μg per injection) or testosterone (T; 300 μg per injection) were injected I.P. two times a week for three weeks into OVX and GdX mice, respectively.

The Prlr^fl/fl^ line was generated as previously described (Brown et al., 2016). Prlr^fl/fl^ line has inverse lox sites; hence, *Cre*-recombination ablates the Prlr gene and activates GFP in targeted cells.

Estrogen was purchased from Sigma-Aldrich (cat: PHR1353-1G) Testosterone was purchased from Sigma-Aldrich (cat: T1875-1G) 4EGI was purchased from Tocris (Minneapolis, MN). Vehicle for IL-6, PGE_2_, PRL and ΔPRL was 0.9% saline or PBS. Vehicle for 4EGI-1 was 0.1% DMSO in 0.9% saline.

Human PRL was generated in an *E.coli* expression system containing plasmid with human PRL (Dr. Goffin; INSERM, Paris). Thus, PRL is fully processed, unmodified (i.e. no glycosylation and phosphorylation) and has molecular weight of ≈23 kDa. The Prlr antagonist, Δ1-9-G129R-hPRL (ΔPRL) (Rouet et al., 2010), which is a modified PRL that binds to and blocks the function of Prlr in rat, mouse and human (Bernichtein et al., 2003) was also synthesised by Dr. Goffin (INSERM, Paris). We and others thoroughly confirmed the specificity of ΔPRL using *in vitro* (Bernichtein et al., 2003; Scotland et al., 2011), and *in vivo* studies (Rouet et al., 2010), including using Prlr KO mice (Belugin et al., 2013).

### RT-PCR

RT-PCR was performed on hindpaw, L3-L5 DRG, and spinal cord (SC) total RNA. Dissected tissue was stored in RNA Later at −20 degrees (Qiagen, Valencia, CA, USA). RNA extraction was done using the QIAzol lysis reagent and the RNAeasy Mini Kit (Qiagen), and manufacturer’s instructions were followed. cDNA was synthesized using Superscript III First Strand Synthesis kit (Invitrogen, CA, California, USA). Primers were: Prlr-F (5’-CCATTCACCTGCTGGTGGAATCCT-3’), GFP-F (5’-AAGGCTACGTCCAGGAGCGCACCA-3’), GFP-R1 (5’-CGTCCTCGATGTTGTGGCGGATC-3’) and GFP-R2 (5’-TGGTGCGCTCCTGGACGTAGCCTT-3’). Amplification of target sequences was detected on 1 or 1.5% agarose gel depending on band size.

### Behavior experiments

Hyperalgesic priming was established using a previously described model (Aley et al., 2000; Kim et al., 2016). IL-6 was injected intra-plantarly (I.Pl.), which created a transient mechanical hypersensitivity and initiated hyperalgesic priming. After IL-6-induced mechanical hypersensitivity resolved and thresholds returned to a baseline level, PGE_2_ was administered either I.Pl. or intrathecally (I.T.) to precipitate the primed state and induce mechanical hypersensitivity. PRL, ΔPRL, or 4EGI-1 were administered immediately prior to IL-6 or PGE_2_ administration. To evaluate mechanical hypersensitivity following the I.Pl. or I.T. injections, animals were habituated for 45-60 minutes in elevated behavior racks and then paw withdraw threshold was determined using the up-down von Frey method (Chaplan, et al., 1994). Both the experimenters performing the behavior and data analysis were done blinded.

### Experimental Design and Statistical Analysis

GraphPad Prism 7.0 (GraphPad, La Jolla, CA) was used for all statistical analyses of data. Data are presented as mean ± standard error of the mean (SEM), with “n” referring to the number of independent animals per group in behavioral experiments. The sex of the animals used in each experiment is described in the text. Differences between groups were assessed by either mixed effects or repeated measures ANOVA with Bonferroni post-hoc tests and is noted for each figure. Statistically significance was determined as p<0.05. Interaction F ratios, and the associated p values are reported in the text.

## Results

### Sex differences in hyperalgesic priming in mice are regulated by gonadal hormones and translation regulation

Chronic pain occurs more frequently in females (Unruh, 1996; Berkley, 1997; Fillingim et al., 2009; Traub and Ji, 2013). This difference could be mediated by sex-dependent mechanisms controlling the transition from acute to chronic pain. We used the hyperalgesic priming paradigm (Aley et al., 2000) to gain insight into female-specific mechanisms involved in the acute to chronic pain transition. In our experiments, hyperalgesic priming was initiated with an intraplantar (I.Pl.) injection of IL-6 (0.5 ng). When the initial hypersensitivity from this IL-6 injection had resolved, the presence of priming was assessed with either an I.Pl. or intrathecal (I.T.) injection of PGE_2_ (0.1 μg; *Figure* 1A). Female C57BL/6 mice that were primed with IL-6 and then subsequently received an I.Pl injection of PGE_2_ had a slightly longer persistence of mechanical hypersensitivity when compared to males (repeated measures ANOVA; F (11, 88) = 1.437; P=0.0.1708; n=5; *Figure 1B*). I.T. injection of PGE_2_ in females had a significant longer duration of response to PGE_2_ when compared to males (repeated measures ANOVA; F (14, 91) = 8.975 P<0.0001 n=5; *Figure 1C*). To rule out a possible strain-dependent sex effect the experiment with spinal administration of PGE_2_ following I.Pl. IL-6 was repeated in male and female Swiss Webster (SW) mice. The difference in the length of the PGE_2_ response following I.T. administration was even longer lasting in this outbred strain (repeated measures ANOVA; F (10, 140) = 8.409; P<0.0001; n=8; *Figure 1D*). To gain insight into the mechanistic underpinnings of the sex difference we identified, we did the remaining experiments with I.Pl. injection of IL-6 and I.T. administration of PGE_2_ in C57BL/6 mice.

**Figure 1:**
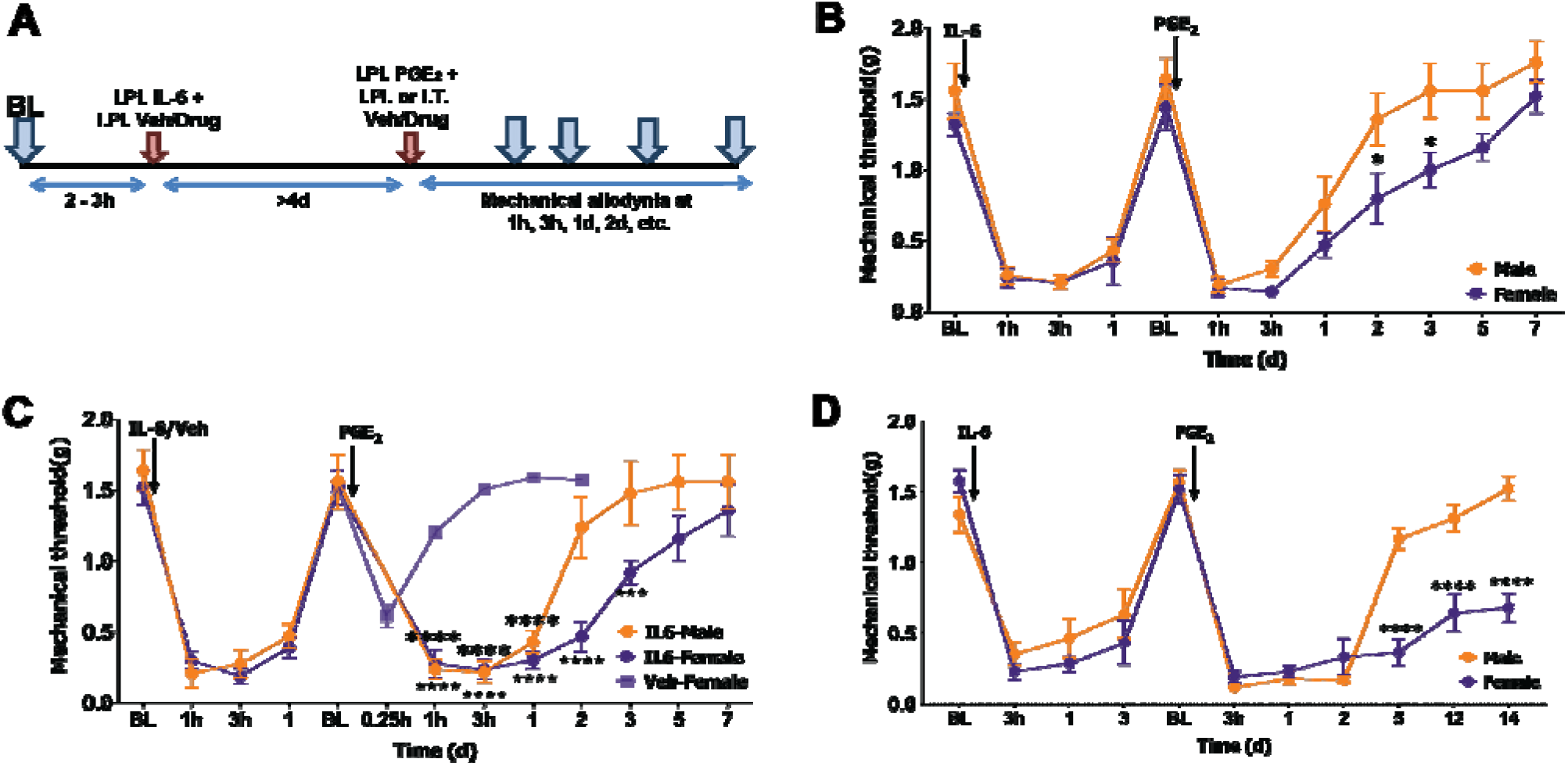
Persistence of hyperalgesic priming is greater in female mice. (**A**) Schematic of the IL-6-induced hyperalgesic priming model. BL, baseline measurements; SC, spinal cord. Brown arrows are injection time points. Blue arrows are post-PGE_2_ mechanical nociception measurement time points. (**B**) Hyperalgesic priming model; IL6 priming into paw and PGE_2_ injection into paw of female and male C57BL/6 mice. (**C**) Hyperalgesic priming model; IL6 priming into paw and PGE_2_ injection into spinal cord of female and male C57BL/6 mice. (**D**) The same model as in *panel C* in female and male ICR mice. Injection time points for IL-6/Veh and PGE_2_ are indicated by arrows. Repeated measures ANOVA with Bonferroni post-hoc test (For Fig 1C, IL-6-male compared to Veh-Female: * = p<0.05; *** = p<0.001; **** = p<0.0001. IL6-Female compated to Veh-Female: ★★★★= p<0.0001; ★★★= p<0.001. For all others * = p<0.05; *** = p<0.001; **** = p<0.0001. n=5-8).

In the above and following experiments the female animals were all in the estrous phase of the estrous cycle at the time of IL-6 injections. The hyperalgesic priming model lasts at least a week, and despite controlling for the estrous phase at the time of IL-6 injection, female mice cycle quickly (4-5 days) through other phases of the cycle and circulating blood estrogen (E-2) levels will vary on a day-to-day basis. Hence, to control for E-2 levels we used ovariectomized (OVX) and OVX with E-2 supplementation (OVX-E-2) female mice. The substantial reduction of circulating E-2 in OVX mice (Green et al., 2016) did not change the initiation phase of priming, but the persistence of the response following the PGE_2_ injection was significantly shorter (Mixed measures ANOVA; F (34, 220) = 8.710 P=0.0011; n=5-6, *Figure 2A*). Administering E-2 to keep circulating E-2 at approximately proestrus phase levels in OVX-E-2 mice (Green et al., 2016) resulted in a significant extension of both the initiation (IL-6) and priming (PGE_2_) phases in comparison to intact females in the hyperalgesic priming model. Testosterone (T) mediates male-specific nociceptive responses (Sorge et al., 2011). Gonadectomized (GdX) mice have almost no circulating T (Green et al., 2016). These GdX mice had a substantially longer response to IL-6 injection than intact male mice. Males with IL-6-induced mechanical hypersensitivity usually return to baseline mechanical sensitivity within 4-5 days (Kim et al., 2016) (*Figures 1B-1D*). In contrast, the initiation phase of hyperalgesic priming lasted >31d in GdX males, and only partially recovered to baseline levels, which we displayed on a separate graph (*Figure 2B*). The PGE_2_ phase was also lengthened in GdX compared to intact male mice (Repeated measures ANOVA; F (12, 84) = 5.016; P<0.0001; n=5-6; *Figure 2C*). Testosterone rescue in GdX-T animals returned the initiation and priming phases to the same timeline as intact males (*Figure 2C*). These results show that the time course of hyperalgesic priming is more pronounced in female mice when compared to male mice, and is closely regulated by circulating gonadal hormones in both sexes.

**Figure 2:**
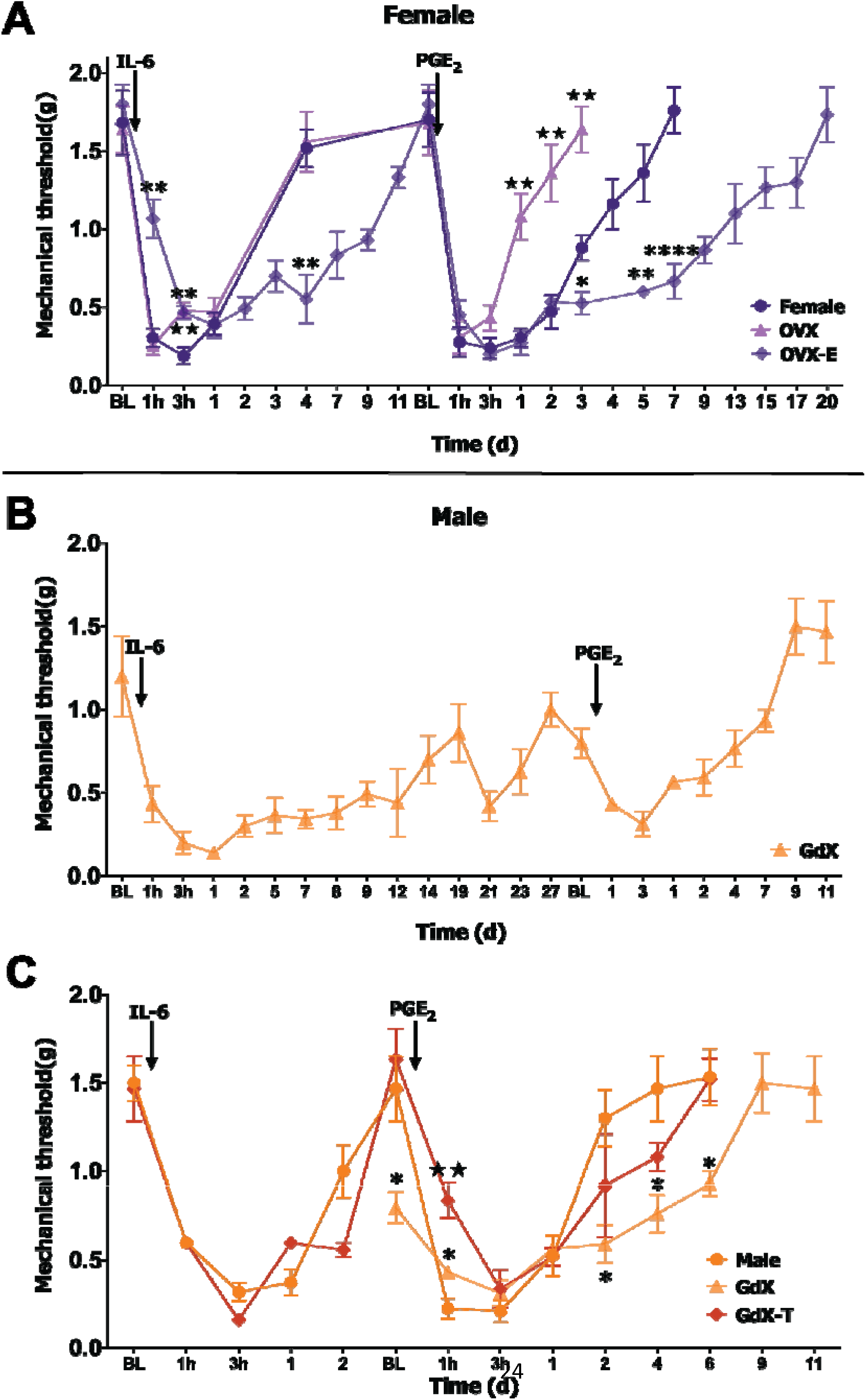
Contribution of gonadal hormones profoundly influence hyperalgesic priming in female and male mice. (**A**) Hyperalgesic priming model with spinal PGE_2_ injection in WT female, OVX and OVX+E (**B**) GdX, and (**C**) WT male, GdX and GdX-T C57BL mice where figure 2B displays the disparity in timelines between GDX animals and those animals that are naïve or GdX-T. Injection time points for IL-6 and PGE_2_ are indicated by arrows. Repeated measures ANOVA with Bonferroni post-hoc test (Fig 1A: *= p<0.05; **= p<0.01; ***= p<0.001; ****= p<0.0001 for OVX-E compared to female and ★★= p<0.01 for OVX compared to female; Fig 1C: *= p<0.05 for GdX compared to Male and ★★= p<0.01 for GdX-T compared to male; n=5-6).

Translation regulation plays a key role in the development of hyperalgesic priming (Price and Inyang, 2015; Khoutorsky and Price, 2018). In previous experiments that were mostly done in male animals, we have shown that inhibition of cap-dependent translation at the time of initiation blocks the development of hyperalgesic priming (Melemedjian et al., 2010; Asiedu et al., 2011; Melemedjian et al., 2014; Moy et al., 2017). Inhibition of cap-dependent translation during the maintenance phase fails to reverse established priming (Asiedu et al., 2011). In concordance with previous experiments (Melemedjian et al., 2010; Asiedu et al., 2011; Melemedjian et al., 2014), the cap-dependent translation inhibitor 4EGI-1 (10 μg) given I.T. immediately prior to PGE_2_ stimulation did not affect PGE_2_-induced mechanical hypersensitivity in males (repeated measures ANOVA; F (8, 64) = 0.3153; P=0.9575; n=5; *Figure 3A*). In stark contrast, 4EGI-1 dramatically reduced the persistence of PGE_2_ precipitated mechanical hypersensitivity in females (repeated measures ANOVA; F (8, 64) = 10.60; P<0.0001; n=5; *Figure 3B*). We next evaluated whether the difference between males and females in the regulation of maintenance of hyperalgesic priming was defined by gonadal hormone status. Removal of circulating E-2 using OVX females abolished the influence of 4EGI-1 on the hyperalgesic priming (repeated measures ANOVA; F (9, 90) = 0.7394; P=0.6719; n=6; *Figure 3C*). These results indicate that the magnitude and maintenance of chronic pain is regulated by gonadal hormones in male and female mice and that the enhanced priming effect seen in intact female mice is dependent on translation regulation at the level of the DRG and/or spinal cord. Interestingly, previous work done entirely in male rodents suggested that translation regulation events at the level of the DRG and/or spinal cord were not involved in maintenance of hyperalgesic priming (Asiedu et al., 2011; Ferrari et al., 2015).

**Figure 3:**
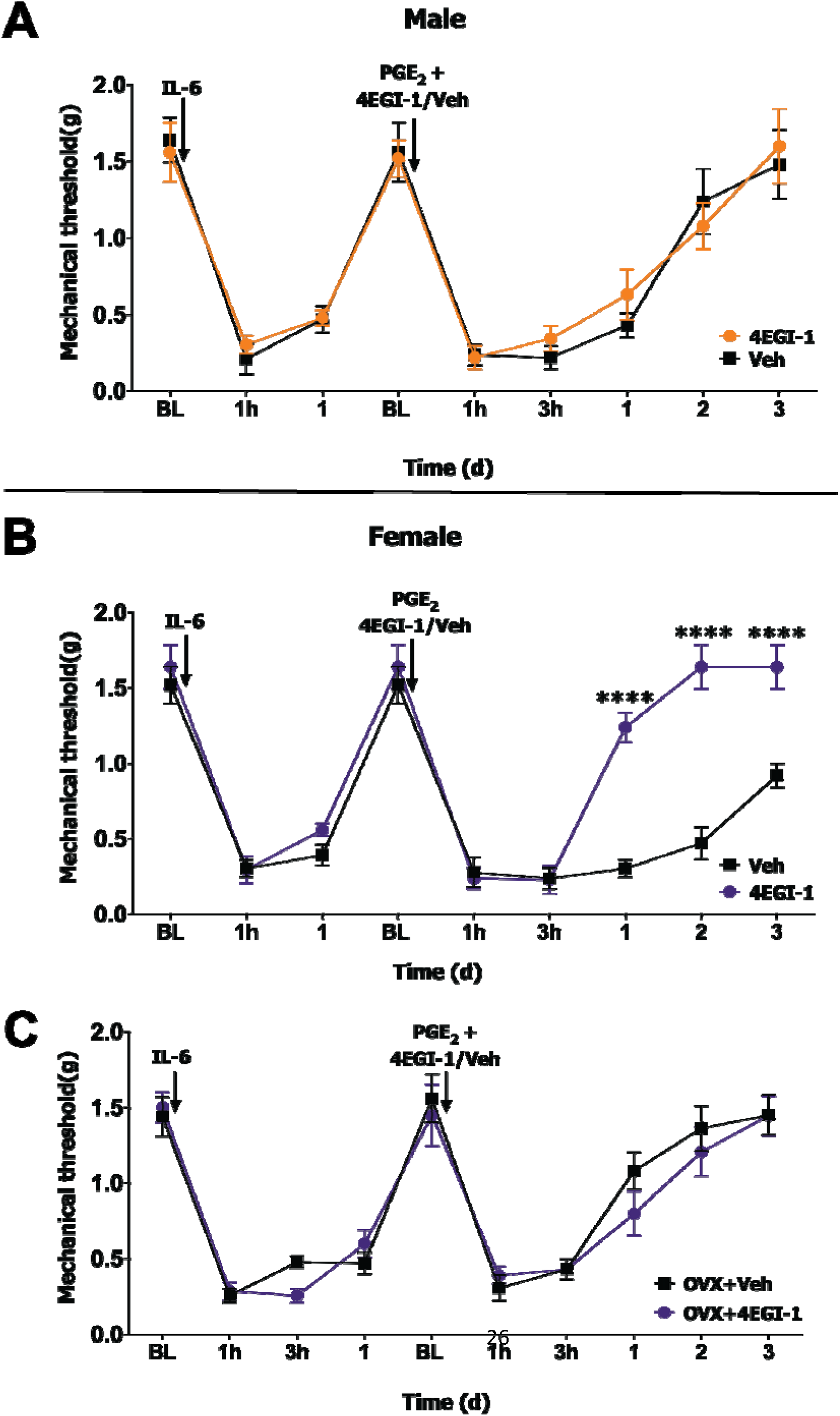
Spinal local translation only contributes to hyperalgesic priming in intact female mice. (**A**) Hyperalgesic priming model with spinal PGE_2_ injection in WT male, (**B**) WT female and (**C**) OVX mice. 4EGI-1 (10μg) or vehicle was administrated spinally at 30 min prior to PGE_2_ injection. Injection time points for IL-6 and 4EGI-1/PGE_2_ are indicated by arrows. Repeated measures ANOVA with Bonferroni post-hoc test (**** p<0.0001; n=5-6).

### A female-specific role for sensory neuronal Prlr in hyperalgesic priming

Responsiveness to PRL in sensory neurons is substantially higher in females (>40 fold) than in males (Patil et al., 2013b; Patil et al., 2019b; Patil et al., 2019a), and strictly controlled by E-2 (Diogenes et al., 2006; Patil et al., 2019b). Endogenous and extra-pituitary PRL is elevated in paw and spinal cord after inflammation and surgical injury (Scotland et al., 2011; Patil et al., 2013a). Accordingly, we examined whether mimicking the presence of endogenous PRL after injury could prolong mechanical hypersensitivity induced by spinal PGE_2_ in both females and males. To do this, we primed the nociceptive pathway with a single injection of exogenous PRL (1μg) I.Pl. (this PRL dosage is active in females but not males (Patil et al., 2019b)) and precipitated hyperalgesic priming with I.T. PGE_2_ at 1d post-PRL treatment (*Figure 4A*). We were able to administer PGE_2_ at 24 h post I.Pl. PRL because 1 μg PRL produces mechanical hypersensitivity for only approximately 3-4 h in females (Patil et al., 2019b). PGE_2_ evoked mechanical hypersensitivity was substantially longer lasting in PRL treated female mice compared to vehicle-primed females (repeated measures ANOVA; F (6, 60) = 6.398; P<0.0001; n=6; *Figure 4C*), but this was not the case in male mice (repeated measures ANOVA; F (4, 40) = 0.2318; P=0.9189; n=5-7; *Figure 4B*).

**Figure 4:**
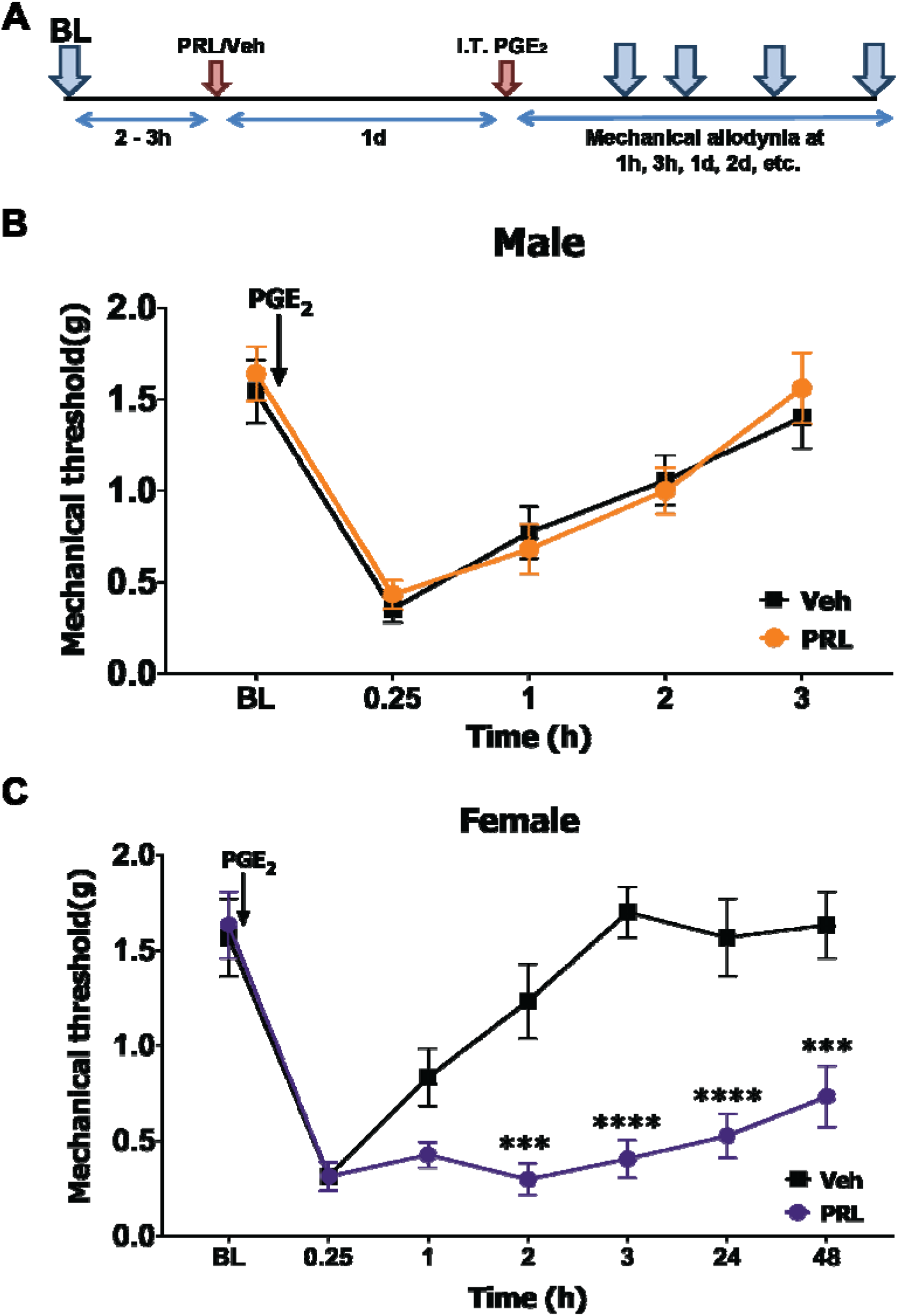
Peripheral PRL only induces hyperalgesic priming in female mice. (**A**) Schematic of the PRL-induced hyperalgesic priming model. BL, baseline measurements after PRL-induced hypersensitivity is fully resolved; SC, spinal cord. Brown arrows are injection time points. Blue arrows are post-PGE_2_ treatment mechanical nociception measurement time points. (**B, C**) Hyperalgesic priming model; PRL or vehicle priming into paw and PGE_2_ injection into spinal cord of male (**B**) or female (**C**) C57BL mice. Injection time points for PGE_2_ are indicated by arrows. Repeated measures ANOVA with Bonferroni post-hoc test (*** p<0.001; **** p<0.0001; n=5-7).

Our present findings suggest that a translation regulation event at the spinal level is critical for enhanced pain chronification in female mice. Our previous work demonstrated that Prlr signaling in the central terminals of nociceptors is important for acute pain models, specifically mechanical hypersensitivity in response to inflammation and injury in females (Patil et al., 2019b). This sex difference can be accounted for by increased translation of *Prlr* mRNA in central terminals of female mice (Patil et al., 2019b) suggesting transport of this mRNA to central terminals of nociceptors. To gain better insight into regulation of *Prlr* mRNA localization, we used *Prlr*^fl/fl^ mice and crossed them with Nav1.8^cre/-^ animals to generate a Nav1.8^cre/-^/*Prlr*^fl/fl^ (*Prlr* CKO) in a set of sensory neurons (*Figure 5A*). The *Prlr*^fl/fl^ line has inverse lox sites; hence, *Cre*-recombination ablates the *Prlr* gene and activates GFP expression in targeted cells driven by the *Prlr* promoter (*Figure 5A*). Analysis of the *Prlr* CKO showed that the truncated transgene *Prlr* mRNA contains the entire 5’-UTR; 4 exons and an intron between exon 1-4 (E1-E4) and GFP, but it does not have the remaining *Prlr* exons or the 3’-UTR (*Figure 5A*). Thus, RT-PCR with *Prlr*-F and GFP-R1 primers (red arrows; *Figure 5A*) produced a 2500 bp band containing an intron sequence from total RNA of female *Prlr* CKO, but not *Prlr*^fl/fl^ (*Figure 5B*). Interestingly, the 2500bp PCR product was detected in spinal cord mRNA suggesting constitutive axonal transport of *Prlr* mRNA (band densities - DRG: 36.06±6.24 vs SC: 123.95±17.12; n=3; *Figure 5B*). To confirm this result, we PCR-amplified GFP (253 bp-band) with GFP-F and GFP-R2 primers (blue arrows; *Figure 5A*) using DRG and spinal cord mRNA from *Prlr*^fl/fl^ and *Prlr* CKO mice. Again, GFP mRNA was detected not only in DRG of *Prlr* CKO females, but also in spinal cord (band densities - DRG: 30.22±5.01 vs SC: 81.58±13.33; n=3; *Figure 5C*). We next examined PGP9.5 mRNA (*Uchl1* gene), which undergoes axonal transport in DRG neurons (Willis et al., 2005; Willis et al., 2007). *Uchl1* mRNA was found in the hindpaw (HP), but at an apparently lower level than observed in the DRG (HP: 25.38±5.58 vs DRG: 110.93±11.84, n=3; *Figure 5D*). In contrast, the construct activated in Prlr CKO containing the Prlr 5’UTR, Prlr E1-E4 and GFP mRNA was at higher levels in the hindpaw than in the DRG (*Figure 5E*). These findings suggest that female *Prlr* mRNA is translocated to sensory neuronal peripheral and central terminals.

**Figure 5:**
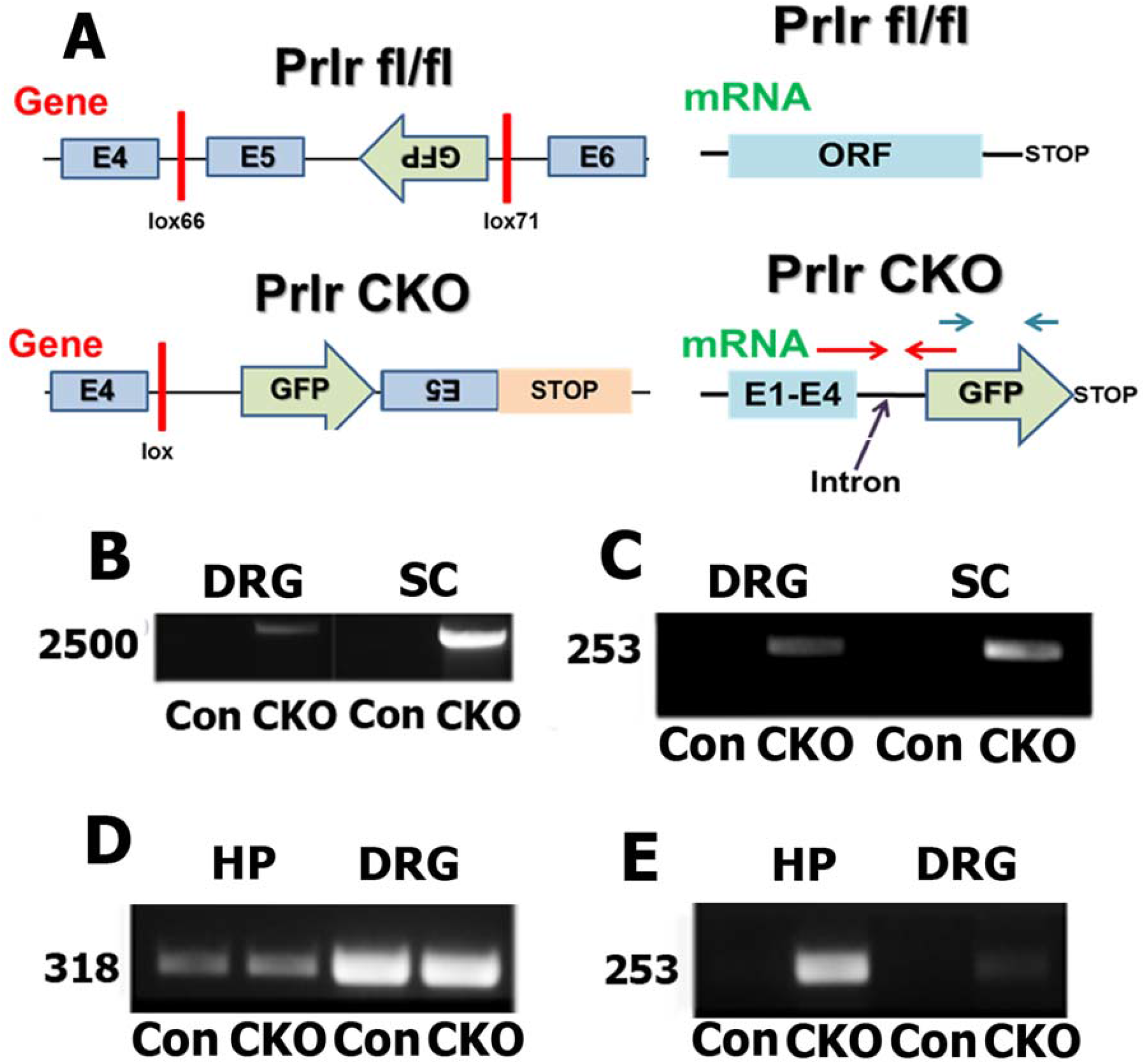
Evidence for translocation of Prlr mRNA to sensory neuronal peripheral and central terminals in female mice. (**A**) Schematic of Prlr^fl/fl^ and Nav1.8^cre/-^/Prlr^fl/fl^ (Prlr CKO) genes and corresponding transcribed mRNA from these genes. Location of Prlr-F and GFP-R1 (red arrows) and GFP-F and GFP-R2 (blue arrow) are shown. (**B**) PCR between Prlr exon 4 and GFP with Prlr-F (exon 4) and GFP-R2 primers for DRG and spinal cord (SC) mRNA from Prlr^fl/fl^ (Con) and Prlr CKO of female mice. (**C**) PCR for GFP with GFP-F and GFP-R1 primers for DRG and SC mRNA from Prlr^fl/fl^ (Con) and Prlr CKO of female mice. PCR for PGP9.5 (**D**) and GFP with GFP-F and GFP-R1 primers (**E**) for DRG and hindpaws (HP) mRNA from Prlr^fl/fl^ (Con) and Prlr CKO female mice.

Next, we used *Prlr* CKO mice to examine the contribution of sensory neuronal Prlr to the regulation of the development of chronic pain in females and males. Prlr ablation in male sensory neurons did not affect either the initiation or persistant phase in the hyperalgesic priming model (repeated measures ANOVA; F (12, 66) = 9.445; P<0.001; n=4-6; *Figure 6A*). In contrast, *Prlr* CKO female mice showed both a significant reduction in mechanical hypersensitivity in response to IL-6 injection and these mice also showed a greatly reduced response to PGE_2_ injection – persistent phase - compared to *Prlr*^fl/fl^ mice (repeated measures ANOVA; F (12, 120)=1.309, P=0.2219; n=5-7; *Figure 6B*). We conclude from this experiment that Prlr in sensory neurons plays a key role in initiation and maintenance of chronic pain in female, but not male, mice.

**Figure 6:**
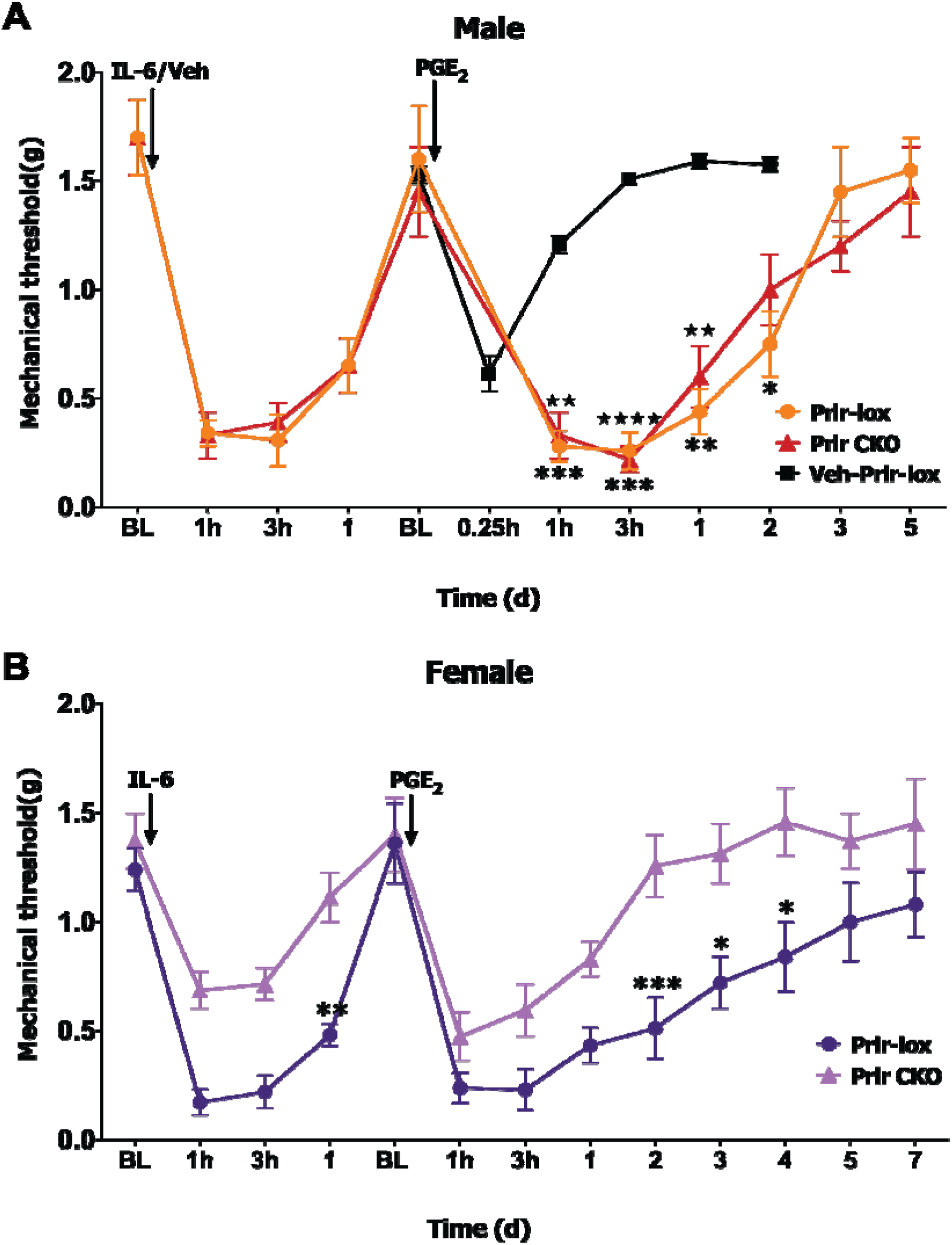
Regulation of hyperalgesic priming by sensory neuronal Prlr selectively in female mice. (**A, B**) Hyperalgesic priming model with peripheral IL-6/Veh and spinal PGE_2_ in Prlr^fl/fl^ (Prlr-lox; control) and Nav1.8^cre/-^/Prlr^fl/fl^ (Prlr CKO) male (**A**) and female (**B**) mice. Injection time points for IL-6/Veh and PGE_2_ are indicated by arrows. Statistic is repeated measures ANOVA with Bonferroni post-hoc test (Figure 6A: *= p<0.05; **= p<0.01, ***= p<0.001 for Prlr-lox compared to Veh-Prlr-lox; and ★ ★ ★ ★= p<0.0001; ★ ★= p<0.01; for Prlr CKO compared to Veh-Prlr-lox; Figire 6B: *= p<0.05; **= p<0.01, ***= p<0.001 n=4-7).

### Prlr signaling and the initiation and maintenance of hyperalgesic priming in female mice

Ablation of Prlr in sensory neurons does not allow for identification of peripheral or central sites driving hyperalgesic priming or the time course of when Prlr signaling occurs during hyperalgesic priming. To explore this in detail, the Prlr antagonist Δ1-9-G129R-hPRL (ΔPRL) (Rouet et al., 2010; Patil et al., 2019b), was delivered at different locations and time points. A single injection of the antagonist ΔPRL (5μg) into the paw immediately prior to the I.Pl. IL-6 priming injection did not affect mechanical hypersensitivity in response to the IL-6 injection (*Figure 7A*). However, there was a significant reduction in mechanical hypersensitivity during the persistent, post PGE_2_ phase in animals that received ΔPRL compared to those that received the vehicle. These ΔPRL treated female mice returned to baseline levels of mechanical sensitivity within 2 days of the PGE_2_ injection (repeated measures ANOVA; F(30, 195) = 6.043; P<0.0001; n=5-6; *Figure 7A*). I.T. administration of ΔPRL (5μg) immediately prior to I.Pl. IL-6 led to a transient reduction in mechanical hypersensitivity for <3h following I.Pl. IL-6 injection (*Figure 7A*). The degree of mechanical hypersensitivity following PGE_2_ injection was almost identical to that observed with I.Pl. administration of ΔPRL. We then evaluated the role of Prlr signaling in the maintenance of hyperalgesic priming with I.Pl. or I.T. administration of ΔPRL (5μg) prior to PGE_2_ administration. Blockage of Prlr signaling in the paw by I.Pl. injection of ΔPRL prior to the injection of PGE_2_ did not influence mechanical hypersensitivity magnitude or duration. In contrast, I.T. administration of ΔPRL coupled with I.T. PGE_2_ administration led to inhibition of mechanical hypersensitivity and a faster return to baseline (repeated measures; F (30, 210) = 7.192; P<0.0001; n=5-6; *Figure 7B*).

**Figure 7:**
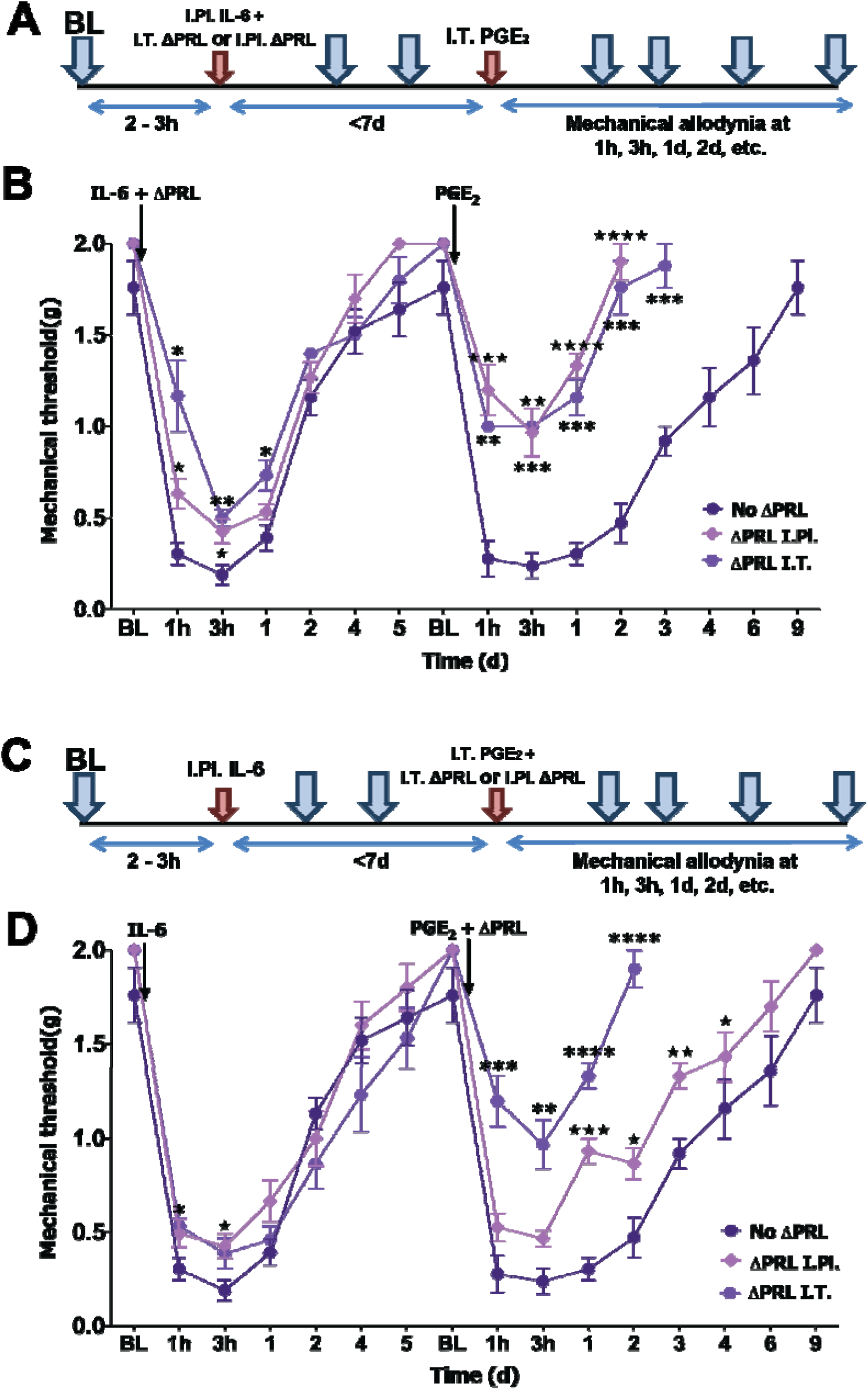
Regulation of the initiation and maintenance of hyperalgesic priming by peripheral and spinal Prlr in female mice. Hyperalgesic priming model in female mice with peripheral IL-6 and spinal PGE_2_. (**A**) Schematic of injection locations and timing for (**B**) Vehicle (No ΔPRL), Prlr antagonist (ΔPRL; 5μg) was co-administrated with IL-6 in paw or spinal cord (SC). (**C**) Schematic of injection locations and timing of (**D**) Vehicle (No ΔPRL) was injected into paw. ΔPRL (5μg) was given into the paw or spinal cord (SC) 30 min before spinal PGE_2_. Repeated measures ANOVA with Bonferroni post-hoc test (*= p<0.05; **= p<0.01; ***= p<0.001; ****= p<0.0001 for ΔPRL I.T. compared to no ΔPRL and ★★★★= p<0.0001; ★ ★★= p<0.001; ★★= p<0.01; ★= p<0.05 for ΔPRL I.Pl. compared to no ΔPRL; n=5-7).

The conventional view is that endogenous PRL comes almost exclusively from the pituitary gland (Ben-Jonathan et al., 2008). However, extra-pituitary sources for PRL have been reported and these PRL sources are especially abundant in humans (BenJonathan et al., 1996). In rodents, inflammation and tissue injury cause an increase in PRL in the paw and spinal cord (Scotland et al., 2011; Patil et al., 2013a). We examined whether endogenous pituitary PRL is involved in the regulation of pain chronicity in male and female mice. Circulating PRL that has originated in the pituitary can be removed by either hypophysectomy (Green et al., 2016) or systemic treatment with bromocriptine (Grattan, 2015). Both approaches have downsides, but we opted to use the systemic bromocriptine approach because bromocriptine is used in clinical studies and hypophysectomy drastically affects gonadal hormone production. We systemically treated both male and female mice with bromocriptine as previously described (Yip et al., 2012). Removal of endogenous pituitary PRL did not influence mechanical hypersensitivity during IL6 phase of hyperalgesic priming initiation in male mice but did slightly prolong the PGE_2_ precipitated hypersensitivity (repeated measures ANOVA; F (10, 90) = 2.980; P=0.0028; n=5-6; *Figure 8A*). In female mice the IL-6 response was enhanced in the bromocriptine treated females but the priming response to PGE_2_ was equivalent in vehicle and bromocriptine treated mice (repeated measures ANOVA; F (14, 126) = 3.127; P=0.0.0003; n=5-6; *Figure 8B*). Collectively, these results suggest that endogenous extra-pituitary PRL signaling plays a key role in hyperalgesic priming in female mice. During initiation of chronic pain this source can be peripheral or central but the crucial source of PRL during the maintenance of pain chronicity is likely central.

**Figure 8:**
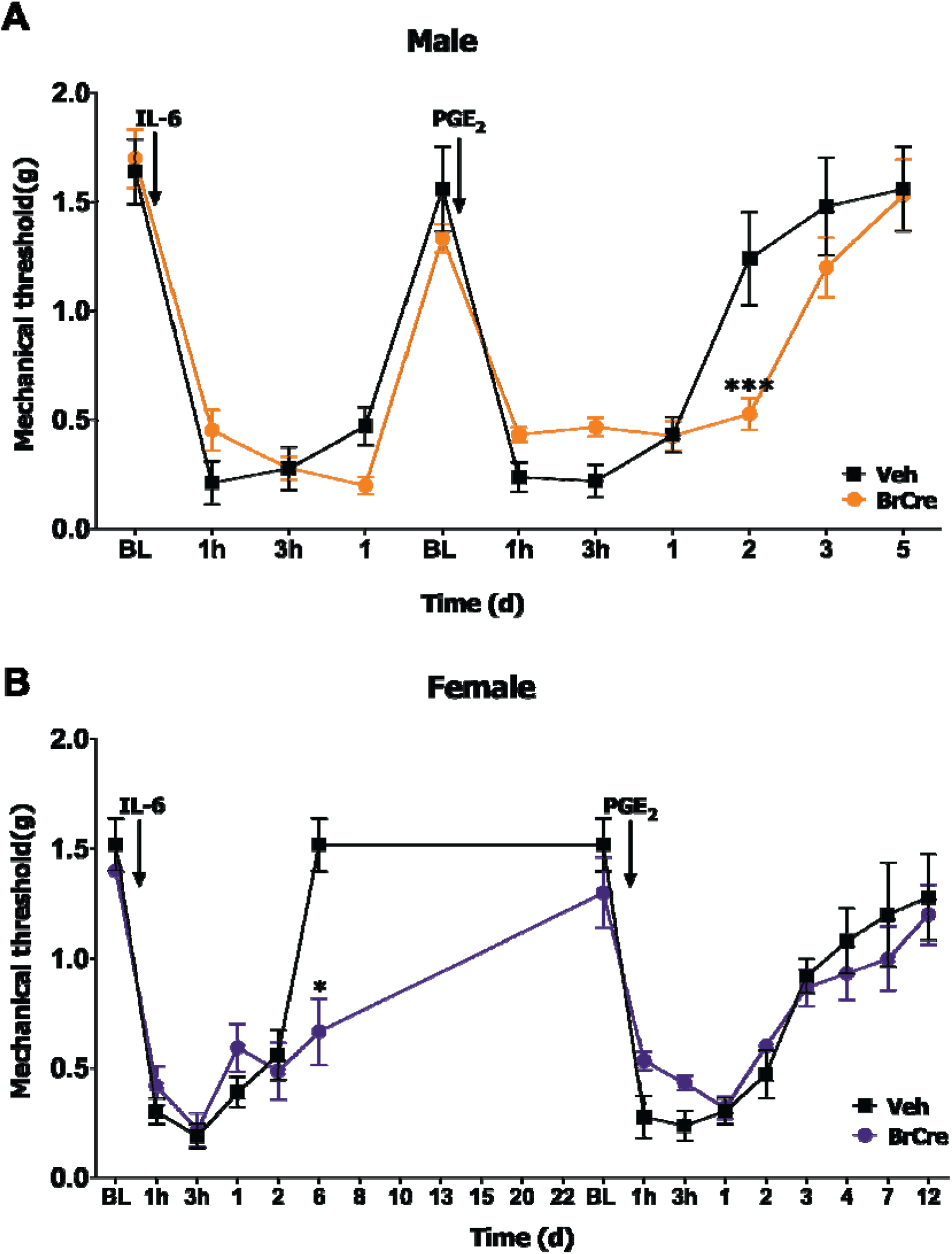
Lack of effect of systemic bromocriptine on hyperalgesic priming in female and male mice. (**A, B**) Hyperalgesic priming model with peripheral IL-6 and spinal PGE_2_ in vehicle and bromocriptine (i.p.; BrCre) treatments of male (**A**) and female (**B**) mice. Injection time points for IL-6 and PGE_2_ are indicated by arrows. Statistic is repeated measures ANOVA with Bonferroni post-hoc test (*= p<0.05, ***= p<0.001; n=5-6).

## Discussion

Studies in both animals and humans demonstrate sexually dimorphic mechanisms controlling the development and resolution of chronic pain (Joseph et al., 2003; Sorge et al., 2015; Nasir et al., 2016; Taves et al., 2016; Lopes et al., 2017; Rosen et al., 2017; Mapplebeck et al., 2018; Paige et al., 2018; Dance, 2019; North et al., 2019; Patil et al., 2019b; Ray et al., 2019a; Rosen et al., 2019). Among these sex differences, several factors have been discovered that drive chronic pain specifically in males (Sorge et al., 2015; Taves et al., 2016; Mapplebeck et al., 2018; Megat et al., 2018; Paige et al., 2018; Shiers et al., 2018; Martin et al., 2019) but relatively little is known about such chronic pain mechanisms in females. There is evidence that these mechanisms are closely regulated by gonadal hormones (Traub and Ji, 2013). For instance, the apparent male-specific effect of microglia-driven P2X4 signaling in neuropathic pain can be conferred to females with testosterone treatment (Sorge et al., 2015). In humans, sex differences in tibial nerve transcriptomes also demonstrate a signature for gonadal hormone influence on sensory neuronal transcriptomes across the lifespan in females (Ray et al., 2019b). Experiments described here clearly demonstrate differential roles of gonadal hormones in development of chronicity in painful conditions with estrogen exacerbating priming effects and testosterone playing a protective role.

Additionally, we demonstrated that translation regulation plays a sex-specific role in the maintenance of chronic pain in female mice. This is especially relevant considering that translation control mechanisms are known contributors to the sensitization of nociceptors (Khoutorsky and Price, 2018; Megat and Price, 2018). Our previous work demonstrated that disruption of translation regulation signaling in the periphery or spinal cord was only capable of interfering with hyperalgesic priming if these treatments were given at the time of the priming event (Melemedjian et al., 2010; Asiedu et al., 2011; Melemedjian et al., 2014). However, these previously published experiments were done entirely in male mice. Our data suggest that targeting these translation regulation mechanisms for the treatment of pain may have additional therapeutic benefits in women. A potential explanation for this differential effect on translation machinery in females is an effect of estrogen on translation machinery (Bronson et al., 2010; Ochnik et al., 2016). E-2-dependent connections between translational control of the Suppressor of Cytokine Signaling (SOCS) and mTOR phosphorylation (Augusto et al., 2010) or regulation of Rheb signaling (Pochynyuk et al., 2006) have been proposed (Matthews et al., 2005; Arbocco et al., 2016). These signaling pathways also play key roles in the excitability of nociceptors (Moy et al., 2017; Khoutorsky and Price, 2018; Megat et al., 2019b; Megat et al., 2019a) and may play a more prominent role in the maintenance of persistent nociceptor plasticity in females than males.

Previous work indicates that sex-dependent mechanisms regulating hypersensitivity in inflammatory and neuropathic pain conditions can be attributed to distinct immune cell types: microglia in males (Sorge et al., 2011; Sorge et al., 2015; Taves et al., 2016; Paige et al., 2018) and T cells in females (Sorge et al., 2015). Importantly, there is an opinion that these cells are regulated by gonadal hormones. Altogether, key molecules involved in sex-dependent regulation of the initiation, maintenance, and resolution would need to (1) be controlled by gonadal hormones, (2) induced by injury, (3) regulate immune cells, (4) undergo local translation control, and (5) be capable of regulating many other genes. The neuroendocrine hormone PRL and its receptor Prlr fit all these requirements. First, Prlr-mediated PRL effects are sex- and gonadal hormone-dependent in many tissues and cell types, including sensory neurons (Torner et al., 2001; Ben-Jonathan et al., 2008; Belugin et al., 2013; Patil et al., 2013a; Patil et al., 2019b; Patil et al., 2019a). Second, many clinical and preclinical studies show that endogenous release of PRL from both pituitary and extra-pituitary origins is induced by inflammation and tissue injury (Chernow et al., 1987; Noreng et al., 1987; BenJonathan et al., 1996; Yardeni et al., 2007; Scotland et al., 2011; Patil et al., 2013a). Third, PRL is an effective direct and/or indirect activator of immune cells, especially macrophages and T-cells (Matera et al., 2001; Savino et al., 2016; Tang et al., 2017). Moreover, many chronic autoimmune diseases affect females more frequently than males, and a potential role for PRL may in part explain this phenomenon for certain autoimmune diseases such as lupus (Tang et al., 2017; Rizzetto et al., 2018). Fourth, translational regulation of Prlr in sensory neurons has been suggested in our previous work (Patil et al., 2019b), and herein we show evidence in support of the translocation of Prlr mRNA to peripheral and central terminals of female or male sensory neurons where it could be translated in a female sex hormone specific fashion (Patil et al., 2019b). Finally, Prlr activation leads to epigenetic changes and transcription regulation of many genes via the STAT5 pathway (Bole-Feysot et al., 1998; Ben-Jonathan et al., 2008). The data presented here is one of the first demonstrations of a female-specific chronic pain initiation and maintenance mechanism acting directly on sensory neurons. Another is calcitonin gene related peptide (CGRP), which is released from sensory neurons but its site of action to produce pain specifically in female mice is not known (Avona et al., 2019). Our findings using a sensory neuron specific knockout of Prlr combined with pharmacological antagonism of Prlr at specific sites suggests that Prlr signaling in sensory neuronal terminals of the spinal cord controls initiation and maintenance of a chronic pain state in female mice. Having said this, we cannot rule out the possible influence of immune cells in our observations, including immune cells as a possible source of PRL that acts on Prlr in the setting of hyperalgesic priming in female mice.

A previous study in rats demonstrated that hyperalgesic priming to carrageenan does not occur in females (Joseph et al., 2003). Subsequent studies in mice and rats have shown additional sexual dimorphisms (Megat et al., 2018; Paige et al., 2018; Inyang et al., 2019), across the lifespan (Moriarty et al., 2019), but none of them have observed a similar absence of priming in female rodents. In fact, our work shows, at least with IL-6 as the priming stimulus, that the magnitude and duration of the response to PGE_2_ given peripherally or intrathecally is longer in female mice than male mice.

In conclusion, our findings demonstrate that sensory neuronal Prlr signaling relies on gonadal hormones and translation mechanisms to contribute to a female-specific regulation of the initiation and maintenance of pain chronicity. These results add a new depth to our understanding of sexually dimorphic signaling pathways involved in chronic pain development. Additionally, our data further substantiate the critical role that the neuroendocrine system and translation regulation play in nociceptor and nociceptive circuit excitability in response to a broad variety of important physiological stimuli.

## Author contribution

C.P. and P.A.B. conducted a majority of experiments, and contributed to manuscript writing and revision; M.P., and J.M. performed certain experiments and analysis; V.G. generated human PRL and Δ1-9-G129R-hPRL, and contributed to experimental design, manuscript writing and revision; D.G. and U.B. generated *Prlr*-cre and Prlr^fl/fl^ mouse lines, and contributed to experimental design, manuscript revision; G.D. helped with experimental design, and contributed to the manuscript preparation; A.N.A. and T.J.P. designed and directed the project, helped with experiments, wrote and edited the first draft of the manuscript; and prepared the final version of the manuscript.

## Acknowledgements

We would like to thank Dr. Dustin Green for advice on the experimental strategy; Dr. Florence Boutillon for producing recombinant PRL and Δ1-9-G129R-hPRL (ΔPRL); This work was supported by NINDS/NIH NS102161 (T.J.P and A.N.A.); NIH/NIGMS GM112747 (A.N.A.); NINDS/NIH NS065926 (T.J.P.); NINDS/NIH NS113457 (CP), and UT BRAIN Pilot Program ID: 1503083 (G.D. and A.N.A.).

## References

Aley KO, Messing RO, Mochly-Rosen D, Levine JD (2000) Chronic hypersensitivity for inflammatory nociceptor sensitization mediated by the epsilon isozyme of protein kinase C. Journal of Neuroscience 20:4680–4685.

Arbocco FCV, Sasso CV, Actis EA, Caron RW, Hapon MB, Jahn GA (2016) Hypothyroidism advances mammary involution in lactating rats through inhibition of PRL signaling and induction of LIF/STAT3 mRNAs. Mol Cell Endocrinol 419:18–28.

Asiedu MN, Tillu DV, Melemedjian OK, Shy A, Sanoja R, Bodell B, Ghosh S, Porreca F, Price TJ (2011) Spinal protein kinase M zeta underlies the maintenance mechanism of persistent nociceptive sensitization. The Journal of neuroscience : the official journal of the Society for Neuroscience 31:6646–6653.

Augusto TM, Bruni-Cardoso A, Damas-Souza DM, Zambuzzi WF, Kuhne F, Lourenco LB, Ferreira CV, Carvalho HF (2010) Oestrogen imprinting causes nuclear changes in epithelial cells and overall inhibition of gene transcription and protein synthesis in rat ventral prostate. Int J Androl 33:675–685.

Avona A, Burgos-Vega C, Burton MD, Akopian AN, Price TJ, Dussor G (2019) Dural Calcitonin Gene-Related Peptide Produces Female-Specific Responses in Rodent Migraine Models. J Neurosci 39:4323–4331.

Belugin S, Diogenes AR, Patil MJ, Ginsburg E, Henry MA, Akopian AN (2013) Mechanisms of Transient Signaling via Short and Long Prolactin Receptor Isoforms in Female and Male Sensory Neurons. J Biol Chem 288:34943–34955.

Ben-Jonathan N, LaPensee CR, LaPensee EW (2008) What can we learn from rodents about prolactin in humans? Endocrine Reviews 29:1–41.

BenJonathan N, Mershon JL, Allen DL, Steinmetz RW (1996) Extrapituitary prolactin: Distribution, regulation, functions, and clinical aspects. Endocrine Reviews 17:639–669.

Berkley KJ (1997) Sex differences in pain. Behav Brain Sci 20:371–380; discussion 435-513.

Bernichtein S, Kayser C, Dillner K, Moulin S, Kopchick JJ, Martial JA, Norstedt G, Isaksson O, Kelly PA, Goffin V (2003) Development of pure prolactin receptor antagonists. J Biol Chem 278:35988–35999.

Bole-Feysot C, Goffin V, Edery M, Binart N, Kelly PA (1998) Prolactin (PRL) and its receptor: actions, signal transduction pathways and phenotypes observed in PRL receptor knockout mice. Endocr Rev 19:225–268.

Bronson MW, Hillenmeyer S, Park RW, Brodsky AS (2010) Estrogen coordinates translation and transcription, revealing a role for NRSF in human breast cancer cells. Mol Endocrinol 24:1120–1135.

Brown RSE, Kokay IC, Phillipps HR, Yip SH, Gustafson P, Wyatt A, Larsen CM, Knowles P, Ladyman SR, LeTissier P, Grattan DR (2016) Conditional Deletion of the Prolactin Receptor Reveals Functional Subpopulations of Dopamine Neurons in the Arcuate Nucleus of the Hypothalamus. The Journal of neuroscience : the official journal of the Society for Neuroscience 36:9173–9185.

Caligioni CS (2009) Assessing reproductive status/stages in mice. Curr Protoc Neurosci Appendix 4:Appendix 4I.

Chernow B, Alexander HR, Smallridge RC, Thompson WR, Cook D, Beardsley D, Fink MP, Lake CR, Fletcher JR (1987) Hormonal responses to graded surgical stress. Arch Intern Med 147:1273–1278.

Childs GV, Unabia G, Miller BT, Collins TJ (1999) Differential expression of gonadotropin and prolactin antigens by GHRH target cells from male and female rats. J Endocrinol 162:177–187.

Dance A (2019) Why the sexes don’t feel pain the same way. Nature 567:448–450.

Diogenes A, Patwardhan AM, Jeske NA, Ruparel NB, Goffin V, Akopian AN, Hargreaves KM (2006) Prolactin modulates TRPV1 in female rat trigeminal sensory neurons. Journal of Neuroscience 26:8126–8136.

Ferrari LF, Araldi D, Levine JD (2015) Distinct terminal and cell body mechanisms in the nociceptor mediate hyperalgesic priming. J Neurosci 35:6107–6116.

Fillingim RB, King CD, Ribeiro-Dasilva MC, Rahim-Williams B, Riley JL (2009) Sex, Gender, and Pain: A Review of Recent Clinical and Experimental Findings. J Pain 10:447–485.

Grattan DR (2015) 60 Years of Neuroendocrinology the Hypothalamo-Prolactin Axis. J Endocrinol 226:T101–T122.

Green DP, Patil MJ, Akopian AN (2016) Influence of hypophysectomy, ovariectomy and gonadectomy on postoperative hypersensitivity in rats. Glob Anesth Perioper Med 2:171–175.

Houghton LA, Lea R, Jackson N, Whorwell PJ (2002) The menstrual cycle affects rectal sensitivity in patients with irritable bowel syndrome but not healthy volunteers. Gut 50:471–474.

Inyang KE, McDougal TA, Ramirez ED, Williams M, Laumet G, Kavelaars A, Heijnen CJ, Burton M, Dussor G, Price TJ (2019) Alleviation of paclitaxel-induced mechanical hypersensitivity and hyperalgesic priming with AMPK activators in male and female mice. Neurobiol Pain 6:100037.

Joseph EK, Parada CA, Levine JD (2003) Hyperalgesic priming in the rat demonstrates marked sexual dimorphism. Pain 105:143–150.

Khoutorsky A, Price TJ (2018) Translational Control Mechanisms in Persistent Pain. Trends Neurosci 41:100–114.

Kim JY, Megat S, Moy JK, Asiedu MN, Mejia GL, Vagner J, Price TJ (2016) Neuroligin 2 regulates spinal GABAergic plasticity in hyperalgesic priming, a model of the transition from acute to chronic pain. Pain 157:1314–1324.

LeResche L, Mancl L, Sherman JJ, Gandara B, Dworkin SF (2003) Changes in temporomandibular pain and other symptoms across the menstrual cycle. Pain 106:253–261.

Liu TT, Qu ZW, Ren C, Gan X, Qiu CY, Hu WP (2016) Prolactin potentiates the activity of acid-sensing ion channels in female rat primary sensory neurons. Neuropharmacology 103:174–182.

Lopes DM, Malek N, Edye M, Jager SB, McMurray S, McMahon SB, Denk F (2017) Sex differences in peripheral not central immune responses to pain-inducing injury. Sci Rep 7:16460.

Mapplebeck JCS, Dalgarno R, Tu Y, Moriarty O, Beggs S, Kwok CHT, Halievski K, Assi S, Mogil JS, Trang T, Salter MW (2018) Microglial P2X4R-evoked pain hypersensitivity is sexually dimorphic in rats. Pain 159:1752–1763.

Martin LJ, Acland EL, Cho C, Gandhi W, Chen D, Corley E, Kadoura B, Levy T, Mirali S, Tohyama S, Khan S, MacIntyre LC, Carlson EN, Schweinhardt P, Mogil JS (2019) Male-Specific Conditioned Pain Hypersensitivity in Mice and Humans. Curr Biol 29:192–201 e194.

Matera L, Mori M, Galetto A (2001) Effect of prolactin on the antigen presenting function of monocyte-derived dendritic cells. Lupus 10:728–734.

Mathew PG, Dun EC, Luo JJ (2013) A Cyclic Pain: The Pathophysiology and Treatment of Menstrual Migraine. Obstet Gynecol Surv 68:130–140.

Matthews J, Almlof T, Kietz S, Leers J, Gustafsson JA (2005) Estrogen receptor-alpha regulates SOCS-3 expression in human breast cancer cells. Biochem Biophys Res Commun 335:168–174.

Megat S, Price TJ (2018) Therapeutic opportunities for pain medicines via targeting of specific translation signaling mechanisms. Neurobiology of Pain:In press.

Megat S, Shiers S, Moy JK, Barragan-Iglesias P, Pradhan G, Seal RP, Dussor G, Price TJ (2018) A Critical Role for Dopamine D5 Receptors in Pain Chronicity in Male Mice. J Neurosci 38:379–397.

Megat S, Ray PR, Tavares-Ferreira D, Moy JK, Sankaranarayanan I, Wanghzou A, Fang Lou T, Barragan-Iglesias P, Campbell ZT, Dussor G, Price TJ (2019a) Differences between dorsal root and trigeminal ganglion nociceptors in mice revealed by translational profiling. J Neurosci.

Megat S, Ray PR, Moy JK, Lou TF, Barragan-Iglesias P, Li Y, Pradhan G, Wanghzou A, Ahmad A, Burton MD, North RY, Dougherty PM, Khoutorsky A, Sonenberg N, Webster KR, Dussor G, Campbell ZT, Price TJ (2019b) Nociceptor Translational Profiling Reveals the Ragulator-Rag GTPase Complex as a Critical Generator of Neuropathic Pain. J Neurosci 39:393–411.

Melemedjian OK, Asiedu MN, Tillu DV, Peebles KA, Yan J, Ertz N, Dussor GO, Price TJ (2010) IL-6- and NGF-Induced Rapid Control of Protein Synthesis and Nociceptive Plasticity via Convergent Signaling to the eIF4F Complex. J Neurosci 30:15113–15123.

Melemedjian OK, Tillu DV, Moy JK, Asiedu MN, Mandell EK, Ghosh S, Dussor G, Price TJ (2014) Local translation and retrograde axonal transport of CREB regulates IL-6-induced nociceptive plasticity. Mol Pain 10:45.

Mogil JS et al. (2011) Pain sensitivity and vasopressin analgesia are mediated by a gene-sex-environment interaction. Nat Neurosci 14:1569–1573.

Moriarty O, Tu YS, Sengar AS, Salter MW, Beggs S, Walker SM (2019) Priming of Adult Incision Response by Early-Life Injury: Neonatal Microglial Inhibition Has Persistent But Sexually Dimorphic Effects in Adult Rats. Journal of Neuroscience 39:3081–3093.

Moy JK, Khoutorsky A, Asiedu MN, Black BJ, Kuhn JL, Barragan-Iglesias P, Megat S, Burton MD, Burgos-Vega CC, Melemedjian OK, Boitano S, Vagner J, Gkogkas CG, Pancrazio JJ, Mogil JS, Dussor G, Sonenberg N, Price TJ (2017) The MNK-eIF4E Signaling Axis Contributes to Injury-Induced Nociceptive Plasticity and the Development of Chronic Pain. J Neurosci 37:7481–7499.

Nasir H, Mahboubi H, Gyawali S, Ding S, Mickeviciute A, Ragavendran JV, Laferriere A, Stochaj U, Coderre TJ (2016) Consistent sex-dependent effects of PKMzeta gene ablation and pharmacological inhibition on the maintenance of referred pain. Mol Pain 12.

Nettleship JE, Jones TH, Channer KS, Jones RD (2007) Physiological testosterone replacement therapy attenuates fatty streak formation and improves high-density lipoprotein cholesterol in the Tfm mouse: an effect that is independent of the classic androgen receptor. Circulation 116:2427–2434.

Noreng MF, Jensen P, Tjellden NU (1987) Per- and postoperative changes in the concentration of serum thyreotropin under general anaesthesia, compared to general anaesthesia with epidural analgesia. Acta Anaesthesiol Scand 31:292–294.

North RY, Li Y, Ray P, Rhines LD, Tatsui CE, Rao G, Johansson CA, Zhang H, Kim YH, Zhang B, Dussor G, Kim TH, Price TJ, Dougherty PM (2019) Electrophysiological and transcriptomic correlates of neuropathic pain in human dorsal root ganglion neurons. Brain 142:1215–1226.

Ochnik AM, Peterson MS, Avdulov SV, Oh AS, Bitterman PB, Yee D (2016) Amplified in Breast Cancer Regulates Transcription and Translation in Breast Cancer Cells. Neoplasia 18:100–110.

Paige C, Maruthy GB, Mejia G, Dussor G, Price T (2018) Spinal Inhibition of P2XR or p38 Signaling Disrupts Hyperalgesic Priming in Male, but not Female, Mice. Neuroscience 385:133–142.

Patil M, Hovhannisyan AH, Wangzhou A, Mecklenburg J, Koek W, Goffin V, Grattan D, Boehm U, Dussor G, Price TJ, Akopian AN (2019a) Prolactin receptor expression in mouse dorsal root ganglia neuronal subtypes is sex-dependent. J Neuroendocrinol 31:e12759.

Patil M, Belugin S, Mecklenburg J, Wangzhou A, Paige C, Barba-Escobedo PA, Boyd JT, Goffin V, Grattan D, Boehm U, Dussor G, Price TJ, Akopian AN (2019b) Prolactin Regulates Pain Responses via a Female-Selective Nociceptor-Specific Mechanism. iScience 20:449–465.

Patil MJ, Green DP, Henry MA, Akopian AN (2013a) Sex-dependent roles of prolactin and prolactin receptor in postoperative pain and hyperalgesia in mice. Neuroscience 253:132–141.

Patil MJ, Ruparel SB, Henry MA, Akopian AN (2013b) Prolactin regulates TRPV1, TRPA1, and TRPM8 in sensory neurons in a sex-dependent manner: Contribution of prolactin receptor to inflammatory pain. American journal of physiology Endocrinology and metabolism 305:E1154–1164.

Pi X, Voogt JL (2002) Sex difference and estrous cycle: expression of prolactin receptor mRNA in rat brain. Brain Res Mol Brain Res 103:130–139.

Pochynyuk O, Medina J, Gamper N, Genth H, Stockand JD, Staruschenko A (2006) Rapid translocation and insertion of the epithelial Na+ channel in response to RhoA signaling. J Biol Chem 281:26520–26527.

Price TJ, Inyang KE (2015) Commonalities between pain and memory mechanisms and their meaning for understanding chronic pain. Prog Mol Biol Transl Sci 131:409–434.

Ray P, Khan J, Wangzhou A, Tavares-Ferreira D, Akopian AN, Dussor G, Price TJ (2019a) Transcriptome analysis of the human tibial nerve identifies sexually dimorphic expression of genes involved in pain, inflammation and neuro-immunity. Frontiers in molecular neuroscience:in press.

Ray PR, Khan J, Wangzhou A, Tavares-Ferreira D, Akopian AN, Dussor G, Price TJ (2019b) Transcriptome Analysis of the Human Tibial Nerve Identifies Sexually Dimorphic Expression of Genes Involved in Pain, Inflammation, and Neuro-Immunity. Front Mol Neurosci 12:37.

Rizzetto L, Fava F, Tuohy KM, Selmi C (2018) Connecting the immune system, systemic chronic inflammation and the gut microbiome: The role of sex. J Autoimmun 92:12–34.

Rosen S, Ham B, Mogil JS (2017) Sex differences in neuroimmunity and pain. J Neurosci Res 95:500–508.

Rosen SF, Ham B, Haichin M, Walters IC, Tohyama S, Sotocinal SG, Mogil JS (2019) Increased pain sensitivity and decreased opioid analgesia in T-cell-deficient mice and implications for sex differences. Pain 160:358–366.

Rouet V, Bogorad RL, Kayser C, Kessal K, Genestie C, Bardier A, Grattan DR, Kelder B, Kopchick JJ, Kelly PA, Goffin V (2010) Local prolactin is a target to prevent expansion of basal/stem cells in prostate tumors. Proc Natl Acad Sci U S A 107:15199–15204.

Savino W, Mendes-da-Cruz DA, Lepletier A, Dardenne M (2016) Hormonal control of T-cell development in health and disease. Nat Rev Endocrinol 12:77–89.

Scotland PE, Patil M, Belugin S, Henry MA, Goffin V, Hargreaves KM, Akopian AN (2011) Endogenous prolactin generated during peripheral inflammation contributes to thermal hyperalgesia. Eur J Neurosci 34:745–754.

Shiers S, Pradhan G, Mwirigi J, Mejia G, Ahmad A, Kroener S, Price T (2018) Neuropathic Pain Creates an Enduring Prefrontal Cortex Dysfunction Corrected by the Type II Diabetic Drug Metformin But Not by Gabapentin. J Neurosci 38:7337–7350.

Slade GD, Bair E, By K, Mulkey F, Baraian C, Rothwell R, Reynolds M, Miller V, Gonzalez Y, Gordon S, Ribeiro-Dasilva M, Lim PF, Greenspan JD, Dubner R, Fillingim RB, Diatchenko L, Maixner W, Dampier D, Knott C, Ohrbach R (2011) Study methods, recruitment, sociodemographic findings, and demographic representativeness in the OPPERA study. J Pain 12:T12–26.

Sorge RE, LaCroix-Fralish ML, Tuttle AH, Sotocinal SG, Austin JS, Ritchie J, Chanda ML, Graham AC, Topham L, Beggs S, Salter MW, Mogil JS (2011) Spinal cord Toll-like receptor 4 mediates inflammatory and neuropathic hypersensitivity in male but not female mice. J Neurosci 31:15450–15454.

Sorge RE et al. (2015) Different immune cells mediate mechanical pain hypersensitivity in male and female mice. Nat Neurosci 18:1081–1083.

Tang MW, Garcia S, Gerlag DM, Tak PP, Reedquist KA (2017) Insight into the Endocrine System and the Immune System: A Review of the Inflammatory Role of Prolactin in Rheumatoid Arthritis and Psoriatic Arthritis. Front Immunol 8:720.

Taves S, Berta T, Liu DL, Gan S, Chen G, Kim YH, Van de Ven T, Laufer S, Ji RR (2016) Spinal inhibition of p38 MAP kinase reduces inflammatory and neuropathic pain in male but not female mice: Sex-dependent microglial signaling in the spinal cord. Brain Behav Immun 55:70–81.

Torner L, Toschi N, Pohlinger A, Landgraf R, Neumann ID (2001) Anxiolytic and anti-stress effects of brain prolactin: improved efficacy of antisense targeting of the prolactin receptor by molecular modeling. J Neurosci 21:3207–3214.

Traub RJ, Ji Y (2013) Sex differences and hormonal modulation of deep tissue pain. Front Neuroendocrinol 34:350–366.

Unruh AM (1996) Gender variations in clinical pain experience. Pain 65:123–167.

Willis D, Li KW, Zheng JQ, Chang JH, Smit AB, Kelly T, Merianda TT, Sylvester J, van Minnen J, Twiss JL (2005) Differential transport and local translation of cytoskeletal, injury-response, and neurodegeneration protein mRNAs in axons. J Neurosci 25:778–791.

Willis DE, van Niekerk EA, Sasaki Y, Mesngon M, Merianda TT, Williams GG, Kendall M, Smith DS, Bassell GJ, Twiss JL (2007) Extracellular stimuli specifically regulate localized levels of individual neuronal mRNAs. J Cell Biol 178:965–980.

Yardeni IZ, Shavit Y, Bessler H, Mayburd E, Grinevich G, Beilin B (2007) Comparison of postoperative pain management techniques on endocrine response to surgery: a randomised controlled trial. Int J Surg 5:239–243.

Yip SH, Eguchi R, Grattan DR, Bunn SJ (2012) Prolactin signalling in the mouse hypothalamus is primarily mediated by signal transducer and activator of transcription factor 5b but not 5a. J Neuroendocrinol 24:1484–1491.

